# Plasticity of interhemispheric motor cortex connectivity induced by brain state-dependent cortico-cortical paired-associative stimulation

**DOI:** 10.1101/2025.04.16.648589

**Authors:** Maria Ermolova, Gábor Kozák, Paolo Belardinelli, Ulf Ziemann

**Affiliations:** Department of Neurology & Stroke, University of Tübingen, Tübingen, Germany; Hertie Institute for Clinical Brain Research, University of Tübingen, Tübingen, Germany; CiMeC, Center for Mind/Brain Sciences, University of Trento, Italy

**Keywords:** Interhemispheric inhibition, functional connectivity, EEG-TMS, brain state-dependent stimulation, plasticity

## Abstract

Transcallosal connectivity between the hand areas of the two primary motor cortices (M1) is important for coordination of unimanual and bimanual hand motor function. Effective connectivity of this M1-M1 pathway can be tested in the form of short-interval interhemispheric inhibition (SIHI) using dual-coil transcranial magnetic stimulation (TMS). Recently, we and others have demonstrated that the phase of the ongoing sensorimotor µ-rhythm has significant impact on corticospinal excitability as measured by motor evoked potential (MEP) amplitude, and repetitive TMS of the high-excitability state (trough of the µ-rhythm) but not other states resulted in long-term potentiation-like MEP increase. Here, we tested to what extent the phase of the ongoing µ-rhythm in the two M1 cortices affects long-term change in SIHI. In healthy subjects we applied cortico-cortical paired associative stimulation (ccPAS) in four different µ-phase conditions in the left conditioning M1 and right test M1 (trough-trough, trough-positive peak, positive peak-trough, random phase). We found long-term strengthening of SIHI but no differential effect of phase conditions. Findings point to a distinct regulation of plasticity of corticospinal vs. M1-M1 connectivity. The observed ccPAS-induced strengthening of effective M1-M1 connectivity (SIHI) may be utilized for therapeutic applications that potentially benefit from modification of interhemispheric excitation/inhibition balance.

## Introduction

Connectivity between the hand areas of the primary motor cortices (M1) in the two hemispheres through the corpus callosum is crucially important for coordination of unimanual and bimanual hand motor function ^1,2^. The transcallosal fibers are glutamatergic but with prevailing projection onto inhibitory interneurons in the receiving M1 ^3^. Effective connectivity of this transcallosal M1-M1 pathway can be tested in the form of short-interval interhemispheric inhibition (SIHI) using dual-coil transcranial magnetic stimulation (TMS), with application of the conditioning stimulus over the hand area of one M1 followed by the test stimulus over the hand area of the other M1 with typical interstimulus intervals of 8-15 ms ^4^.

After lesion, such as a motor stroke, imbalance in transcallosal inhibition, with often exaggerated SIHI from the contralesional to ipsilesional M1 may contribute to persisting motor deficits ^5,6^. Modulation of abnormal transcallosal inhibition may constitute a therapeutic target in supporting stroke neurorehabilitation ^7,8^. One approach towards modulation of M1-M1 connectivity is cortico-cortical paired-associative stimulation (ccPAS) ^9,10^. Several studies consistently demonstrated a reduction of SIHI from the conditioning to test M1 when paired pulses were repetitively applied with an interstimulus interval of 8 ms while an interval of 1 ms or random interval stimulation had no effect ^11,12^. Those and other studies also showed that corticospinal excitability of the test motor cortex, as measured by motor evoked potential (MEP) amplitude increase after M1-M1 ccPAS, with interstimulus intervals of 6-15 ms (i.e., an interval effective to elicit SIHI) ^11,13,14^.

Recently, we have introduced brain state-dependent stimulation approaches, using real-time analysis of electroencephalography (EEG) signals. We demonstrated with this real-time EEG-TMS technology that the induction of long-term potentiation-like change of M1 excitability with repetitive TMS was more effective when consistently targeting an M1 high-excitability state, i.e., the trough of the ongoing sensorimotor µ-rhythm, in comparison to targeting a low-excitability state (i.e., the positive peak of the µ-rhythm) or random phase stimulation ^15,16^.

Here we sought to test, for the first time, brain state-dependent ccPAS, using the real-time EEG-TMS approach. We have demonstrated previously that M1-M1 effective connectivity (i.e., SIHI) is stronger when both M1 are in synchrony with the phase of their ongoing µ-rhythms, compared to out of synchrony conditions ^17^. Based on these previous findings, we hypothesized that brain state-dependent ccPAS will most effectively change SIHI in conditions of high M1-M1 effective connectivity (i.e., when the µ-rhythm of both M1 is in synchrony) and/or a high-excitability state of the conditioning M1 or the test M1 (i.e., when the µ-rhythm in the conditioning or test M1 is at trough). Towards this end, we tested the following four real-time EEG-TMS ccPAS conditions in a blinded randomized crossover design in healthy participants: µ-rhythm of conditioning M1 - test M1 in (1) trough - trough, (2) positive peak - trough, (3) trough - positive peak, (4) random phase stimulation. We assessed the following outcomes 0, 30 and 60 min after ccPAS in comparison to baseline: MEP amplitude elicited from conditioning and test motor cortex, SIHI, and spontaneous functional M1-M1 connectivity as measured by the weighted phase lag index (wPLI) ^18^ between the source signals reconstructed from the resting-state EEG. In addition, to determine whether the phase lag itself is hardwired or susceptible to phase-dependent ccPAS modulation, we directly analyzed phase lag distributions in the same signals pre- and post-intervention ^19,20^.

## Methods

The study protocol was approved by the local ethics committee of the Medical Faculty of the University of Tübingen (protocol 810/BO20212). The procedures conformed to the Declaration of Helsinki and adhered to the current TMS safety guidelines . All participants provided written informed consent before participation.

### Participants

Potential participants (healthy adults ≥18 years, no previous or present neurological or psychiatric comorbidity, no alcohol or drug abuse) were screened for resting motor threshold (RMT) <72% of maximum stimulator output (MSO). RMT was defined as the lowest stimulation intensity eliciting MEP with a peak-to-peak amplitude of at least 50 µV in 50% of trials (Rossini et al., 2015). This inclusion criterion was introduced because higher RMT would have exceeded the possible range of intensities needed for testing the SIHI input-output (IO) curve (see below). Twenty-three participants fulfilling the RMT criterion underwent an anatomical MRI scan to enable navigated TMS, including T1- and T2-weighted images. The final retained sample included 16 participants (10 female; mean age ± SD, 25 ± 4 years). Five subjects had to be excluded because no SIHI was detected in the SIHI IO-curve, one subject was not able to stay alert during the experiments, and one subject withdrew from participation.

### Data acquisition

Scalp EEG was recorded using a TMS compatible Ag/AgCl sintered ring electrode cap (EasyCap, Wörthsee, Germany) with 64 channels in the International 10–20 EEG system arrangement, EMG was recorded using ECG electrodes (CardinalHealth, Dublin, Ireland) in a belly-tendon montage ^21^. Electrodes were placed on the muscle belly of the first dorsal interosseus (FDI) and the proximal interphalangeal joint of the index finger, and the muscle belly of the abductor pollicis brevis (APB) and the thumb’s interphalangeal joint, bilaterally.

A 24-bit, 80-channel biosignal amplifier was used for the EEG and EMG recordings (NeurOne Tesla, Bittium Biosignals) with a sampling rate of 5 kHz. The stimulation setup consisted of two actively cooled TMS figure-of-eight coils (Cool-B35 HO, MagVenture, Farum, Denmark) and a ‘MagPro R30’ and ‘MagPro X100’ stimulator (MagVenture, Farum, Denmark). Both stimulators delivered a biphasic pulse with posterior-to-anterior main direction of the induced current over the hand knob of the right and left M1. The coil location and orientation were identified in a motor hot spot search as eliciting the highest MEP amplitude in the FDI of the contralateral hand ^21^. The participants were seated in a comfortable reclining chair with a neck rest (MagVenture, Farum, Denmark). A vacuum pillow (MagVenture, Farum, Denmark) and two fixation arms (Super Flex Arm, MagVenture, Farum, Denmark) were used to immobilize the head of the subjects and maintain fixed coil positions throughout the experiment. A stereoscopic neuronavigation system (Localite GmbH, Sankt Augustin, Germany) was used to co-register the head of the participants to their individual T1- weighted MRI scan, and to record the position of the EEG electrodes and TMS coils relative to the head of the participants.

### Real-time algorithm

The sensorimotor µ-rhythm was extracted by computing a surface Laplacian montage centered over the C3 and C4 electrodes of the left and right sensorimotor cortex, respectively ^15^. Instantaneous phase was estimated in real-time using a custom-built digital biosignal processor with an algorithm implemented in Simulink Real-Time (Mathworks Ltd, USA, R2017a), for details see previous publications ^15,17^.

### Experimental procedures

The experiment was designed to investigate the long-term effects of cortico-cortical paired associative stimulation (ccPAS) on connectivity between the M1 in the two hemispheres. Connectivity was probed immediately before (baseline) and at 0, 30, and 60 min post-ccPAS, testing effective (causal) M1-M1 connectivity with SIHI IO-curves, and functional M1-M1 connectivity with resting-state EEG.

### SIHI IO-curve and resting-state EEG acquisition

A SIHI IO-curve was obtained in the following way: conditioning stimuli of varying intensity (90% to 140% RMT, in steps of 10% RMT, 15 pulses per intensity), followed by test stimuli (interstimulus interval 10 ms, 120% RMT), were delivered in randomized order, and independently of the ongoing µ-oscillations. Additionally, test stimuli alone, without the preceding conditioning pulse, were randomly intermixed to acquire a baseline of unconditioned MEPs. After each SIHI IO-curve measurement, resting-state EEG recording was made with eyes open (5 min). An additional 10-min resting-state EEG recording was obtained at the beginning of the experimental session, yielding two pre-ccPAS and three post-ccPAS EEG measurements per experimental session.

### ccPAS intervention

The ccPAS intervention consisted of a block of paired-pulse stimuli (a total of 180 pairs, interstimulus interval 10 ms, minimum inter-trial interval 1.5 s) during rest. Whenever the pre-set phase condition of the ongoing µ-oscillation in the left and right sensorimotor cortex was met, a paired-pulse stimulus was triggered with a suprathreshold conditioning stimulus (intensity was set to the inflection point of the baseline SIHI IO-curve) delivered to the left M1, followed by a suprathreshold test stimulus (intensity: 120% of RMT) delivered to the right M1. The trigger condition in a given experimental session was one of the following combinations of positive peak and trough of the ongoing μ-rhythm in the conditioning left and the test right motor cortex: trough-trough, trough-positive peak, positive peak-trough (Fig. 1A). A random phase condition served as control, where the phase of the stimulation for each trigger was determined by the `rand` function of MATLAB. Each participant underwent all four intervention types, each type in a separate experimental session. The order of the interventions was counterbalanced across participants. Sessions of each participant were spaced at least 1 week apart to prevent carryover effects. Participants were blinded to the experimental condition.

**Figure 1.**
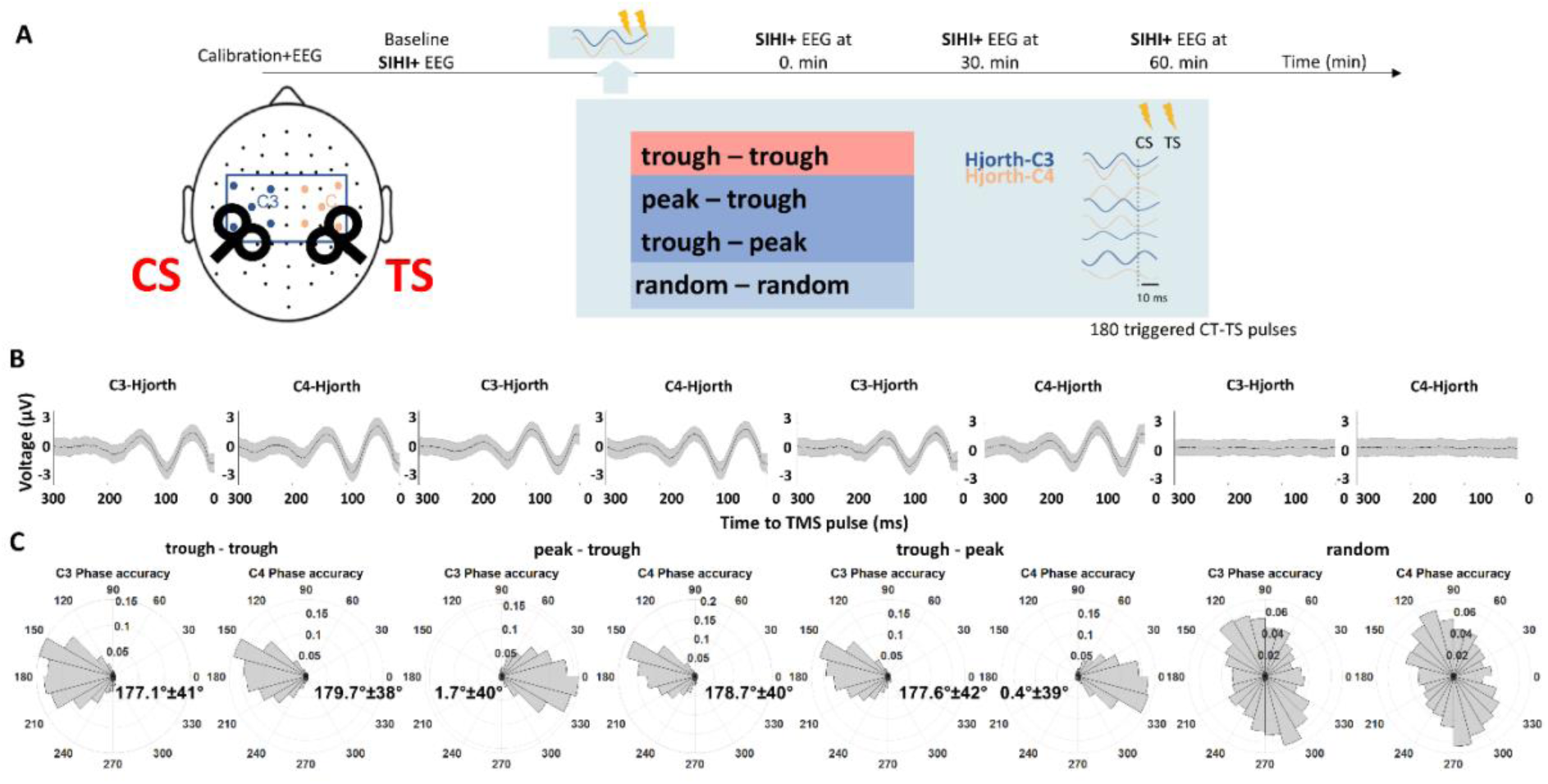
Overview of the methodology. **(A)** Visual summary of the experimental protocol. For detailed description, see experimental procedures. CS, conditioning stimulation; TS, test stimulation. **(B)** Grand mean of the raw Hjorth-C3 and Hjorth-C4 EEG signals across all subjects and trials in each condition preceding the TMS pulse at 0 ms. Shaded area represents ±SEM. **(C)** Cumulative binned distribution of the targeted phase angle at the time of the stimulation for the conditioning motor cortex (left M1, Hjorth-C3) and for the test motor cortex (right M1, Hjorth-C4) in each trigger condition on the population level (n = 16). Phase angles are binned (30°) and probabilities are indicated. Angular means ±1 SD are displayed for all phase-specific conditions.

### EMG preprocessing

EMG data was epoched around the TMS pulse in a time window of [-500 ms 500 ms] using EEGLAB toolbox (v.2024.2, ^22^). Slow drifts were eliminated by Laplacian trendline fitting. After filtering out line noise the TMS-related decay artifact was then removed from the EMG signal by fitting an exponential curve to the signal in each EMG channel and trial separately (for details, see ^23^). Finally, the peak-to-peak MEP amplitude was extracted in a time window of [20 ms 40 ms] after the TMS pulse. Single-trial SIHI values were calculated by dividing each single-trial conditioned MEP amplitude by the mean unconditioned MEP within the same recording. Single-trial SIHI values from the FDI muscle of the left hand (i.e., the test hand contralateral to the right M1 where the test TMS pulses were applied) were used for SIHI analysis.

### EEG preprocessing

EEG data were preprocessed within MATLAB R2023b via EEGLAB toolbox (v.2024.2). The preprocessing was performed in a semi-automatic pipeline, making use of EEGLAB’s automatic preprocessing functions. Data were downsampled to 1000 Hz, re-referenced to the average reference, and highpass-filtered at 1 Hz (high-pass FIR filter, transition band: 0.75–1.25 Hz) to remove slow trends. Bad channels (flat signal, noisy, or affected by line noise) and noisy time segments were identified and removed through EEGLAB’s automatic preprocessing (the parameters were adjusted between datasets as necessary, retaining a minimum of 56 channels and 3 minutes of signal). Oculographic artifacts were removed with ICA (Picard algorithm). EEG data from the same session were concatenated during preprocessing to ensure consistent channel and component removal. After preprocessing, the 10-min recordings were truncated to 5 min to match the maximal duration of the other recordings.

### Source localization

Anatomical T1-weighted MR images were obtained for each participant with a 3T Siemens Prisma scanner. The MR images were used for TMS neuronavigation, for recording EEG electrode positions, and for EEG forward models. A three-shell Boundary Element Model was generated from the segmentation of each participant’s T1-weighted MR images with customized scripts. The source space, as the “midthickness” surface between grey matter and white matter boundaries, was individually created with each of the (roughly) 16K points adjusted for sulci and gyri on a spherical template to preserve comparability across subjects.

To enhance spatial precision, EEG sensor positions from each measurement session were aligned to the scalp mesh using head fiducials. Individual leadfields were then created for each subject and session making use of the pinpointed EEG positions and assuming fixed dipole orientations normal to the cortical surface. Leadfield and EEG signals were referenced to the average reference.

For the selection of dipoles from the sensorimotor hand knobs in the two hemispheres, we defined on our midthickness mesh the union of the three sensorimotor Glasser parcels (1. BA4 (M1) 2. S1, 3. Area 1 ^24^) and the union of the six Brodmann hand areas (BA Hand 1-6). Then, we considered the intersection between the two previous macro-parcels strictly defined on the same mesh at 16K points. In this way, we were able to consider all the sensorimotor areas related to the hand knob in both hemispheres.

Source activity was reconstructed using Linearly Constrained Minimum Variance beamformer ^25^. The covariance matrix used to create the filter was regularized by Tikhonov regularization with a parameter set to 10 times the maximum sensor power^26^. Source signals were averaged across the dipoles of interest within each hemisphere.

### Functional connectivity analysis

Connectivity and phase lag analysis were performed in Python (v.3.12.5) using MNE-Python (v.1.8.0), MNE-connectivity (v.0.7.0), Numpy (v.1.26.4), Scipy (v.1.14.1), and Pandas (v.2.2.2) toolboxes. Source signals were downsampled to 200 Hz and epoched into 2-s windows with 75% overlap. Time-frequency decomposition was done in the 8-13 Hz range using Morlet wavelets with the length of 5 cycles. Weighted Phase Lag Index (wPLI) ^18^ was calculated across time and averaged across frequencies within each epoch.

### Phase lag distribution analysis

Source signals were downsampled to 200 Hz, filtered in the 8-13 Hz range (band-pass, zero-phase, non-causal FIR filter, Hamming windowed, lower and upper -6 dB cut-off frequencies: 7 Hz and 14.62 Hz, filter length: 331 samples), further down-sampled to 50 Hz to facilitate computational speed, and transformed into analytic signals with the Hilbert transform. Phase lags were calculated for each time sample, unwrapped to prevent discontinuities exceeding π between consecutive time samples, and rotated to span 0 to 2π.

Phase lag samples were selected for analysis such that they coincided with high power and high connectivity states. For that purpose, data were epoched into 2-s non-overlapping epochs. Connectivity threshold was determined for each recording as a median wPLI value across the epochs. Power threshold was derived as a median envelope amplitude across the epochs. Phase lags from epochs whose wPLI and median power both exceeded threshold were used in the subsequent analysis.

### Statistical analyses

Statistical analyses were conducted either in MATLAB using Statistics and Machine Learning Toolbox and Circular Statistics Toolbox (Berens, 2009) or in R (v.4.4.1) using lme4 (v.1.1-35.5, for model selection), lmerTest (v.3.1-3, for the final model fit), performance (v.0.12.4, for evaluation of model residuals), and emmeans (v.1.10.5, for post hoc comparisons) packages. Circular statistical analysis was performed with the circular package (v.0.5-1).

### SIHI analysis

A linear mixed-effects (LME) modeling evaluated the effects of ccPAS intervention, time, stimulation intensity, and their interactions on SIHI, accounting for random variability across participants and sessions. SIHI values were transformed using a power of 0.25 (*SIHI*^0.25^) to approximate normal distribution of residuals in the LME model. An LME model was fitted with transformed single-trial SIHI values as a dependent variable and **Intervention Type, Time, Stimulation Intensity**, and their interactions as fixed effects. **Intervention** represented which relative phase was used during ccPAS intervention (4 levels: trough-trough, trough-positive peak, positive peak-trough, random phase). **Time** represented at which time with respect to the intervention was the testing performed (4 levels: baseline, 0, 30 and 60 min post-intervention). **Intensity** represented the intensity of the conditioning TMS pulse delivered during the testing (spanning across 90-140% RMT in 10% RMT steps). **Intervention** and **Time** were modeled as categorical factors, while **Intensity** was treated as a continuous factor. We compared two models with two different random effects structures to see which one will give the best fit. One model included **Subject** as a random intercept while the other model additionally included **Session** nested within **Subject**. The models were compared using Akaike Information Criterion (AIC) and Bayesian Information Criterion (BIC). The final model was recomputed again with restricted maximum likelihood estimation to test for significance of the fixed effects. The p-values were derived via Type II Wald Chi-square Analysis-of-Variance table. Model residuals were visually assessed for normality, homoscedasticity, and linearity. Post hoc comparisons were performed on estimated marginal means, and the p-values were adjusted for multiple comparisons with false discovery rate (FDR) correction. Post hoc pairwise comparisons of **Time** levels tested post-intervention points against pre-intervention baseline. Comparisons of **Intervention** levels tested all other intervention conditions against the random-phase condition. Given significant interactions between factors, pairwise comparisons of one factor were nested in the levels of the interacting factor.

### Analysis of corticospinal excitability

We evaluated the effects of ccPAS intervention, time and stimulation intensity on corticospinal excitability, as measured by MEP amplitude. Separate LME models were computed for the conditioning and test M1, and for both conditioned and unconditioned probing stimulation in case of the test M1. For the conditioning M1, only data from paired-pulse trials (i.e., involving the conditioning pulse) were analysed. The analysis pipeline was identical to that of SIHI, with two exceptions. For MEPs recorded from the right hand FDI in response to conditioning pulses, subthreshold stimulation (90% RMT) was excluded to maintain the linear fit of stimulation intensity as a predicting factor. For MEPs recorded from the left hand FDI in response to unconditioned stimulation, intensity of the test pulse was fixed to 120% RMT as per the experimental protocol and was therefore not included as a factor in the statistical model.

### Connectivity analysis

We tested whether phase-dependent ccPAS changed the strength of resting-state phase-coupling between the two M1 regions, estimated with wPLI. LME analysis evaluated the effect of intervention, time, as well as interactions between them, on wPLI. WPLI values were transformed into the dependent variable by taking the inverse hyperbolic tangent (*atanh*) and subsequently using a power of 0.24 (*atanh*(*wPLI*)^0.24^), to approximate normal distribution of residuals in the LME model. The LME model was computed with transformed wPLI values as a dependent variable and **Intervention**, **Time**, and their interaction as fixed effects. All effects were modeled as categorical factors. The rest of the analysis followed the same procedure as for SIHI (described above). When comparing the levels of **Time**, pre- and post-intervention time points were compared to the second pre-intervention baseline.

### Correlation analysis

We investigated whether changes in corticospinal excitability in the left and right M1 were associated with changes in SIHI by computing two-sided Pearson’s correlations between SIHI vs. MEP amplitudes from the conditioning M1 and test M1. Transformed response variables, as used in the respective LME models, were averaged across trials, intervention conditions, and, where applicable, stimulation intensities. The resulting values were z-score normalized across time points within each subject. The same analysis was conducted to assess the relationship between effective (SIHI) and functional (wPLI) connectivity. For wPLI, the first baseline time point was excluded prior to normalization, as it was not available for SIHI.

### Phase lag analysis

We investigated whether phase-dependent ccPAS altered the phase shift between the two M1 regions. Specifically, we tested whether the distribution of phase lags shifted post-intervention compared to pre-intervention in different directions depending on the type of **Intervention**. Circular mean phase lag was computed for pulled pre-intervention and post-intervention measurements, separately for each experimental session and each participant. A difference of mean pre- and post-intervention phase lags was then taken. Circular analysis of variance (ANOVA) evaluated the effect of **Intervention** on the post-pre difference in the mean phase lag across subjects, with significance tested via a log-likelihood ratio.

Additionally, we examined whether the phase lag value was stable on the population level or whether it varied between individuals. Circular ANOVA assessed whether distribution of phase lags, irrespective of **Time** or **Intervention**, differed across **Subjects**.

## Results

### Brain state-dependent ccPAS

Fig. 1B shows the pooled mean C3- and C4-Hjorth prestimulus EEG signal of each phase condition. Note that the trough and positive peaks of the ongoing µ-oscillation were accurately targeted in the corresponding conditions, while the averaged EEG showed a flat line in the random phase condition. Fig. 1C shows the accuracy in real-time phase detection. The phase accuracy of the triggers was evaluated offline, utilizing the pre-stimulus EEG data [-1500 -5] ms before the stimuli. The rose plots in Fig. 1C depict the binned distribution (30° per bin) of targeted phase over all subjects (n = 16), and angular means ±1 SD. The phase accuracy was comparable to previous studies ^15,17^. The random condition was uniformly distributed around the circle (Hodges-Ajne test, *p* = 0.85 for both hemispheres). The mean intertrial interval (± SD) was 4.67 s ± 1.44 s without a difference of mean intertrial intervals between the different conditions (one-way repeated measure ANOVA *F*(3,45) = 1.698, *p* = 0.18).

### Corticospinal excitability

Corticospinal excitability in the left conditioning M1: We found a significant interaction between **Intervention** and **Time** (χ² = 95, df = 9, p < 0.001, Fig. 1S), as well as between **Intervention** and **Intensity** (χ² = 28, df = 3, p < 0.001). Main effects were significant for **Intensity** (χ² = 19,986, df = 1, p < 0.001) and **Time** (χ² = 148, df = 3, p < 0.001). Across all interventions, MEP amplitude increased post-intervention, with the timing of these changes varying by condition (see Table 1S and Fig. 1S for pairwise comparisons of marginal means). No significant differences were found between **Interventions** at any specific time point.

Unconditioned corticospinal excitability in the right test M1: A significant interaction was observed between **Intervention** and **Time** (χ² = 68, df = 9, p < 0.001, Fig. 2S), with the direction and timing of change depending on the intervention type (see Table 2S and Fig. 2S for pairwise comparisons). After trough-trough ccPAS, MEP amplitude initially decreased at 0 min post-intervention but increased at 30 min compared to baseline. In contrast, both trough-positive peak ccPAS and positive peak-trough ccPAS led to a decrease in MEP amplitude, occurring at 60 min and 30 min, respectively. After random phase ccPAS, MEP amplitude increased at 0 min post-intervention. No significant differences were found between **Interventions** at any specific time point.

**Figure 2.**
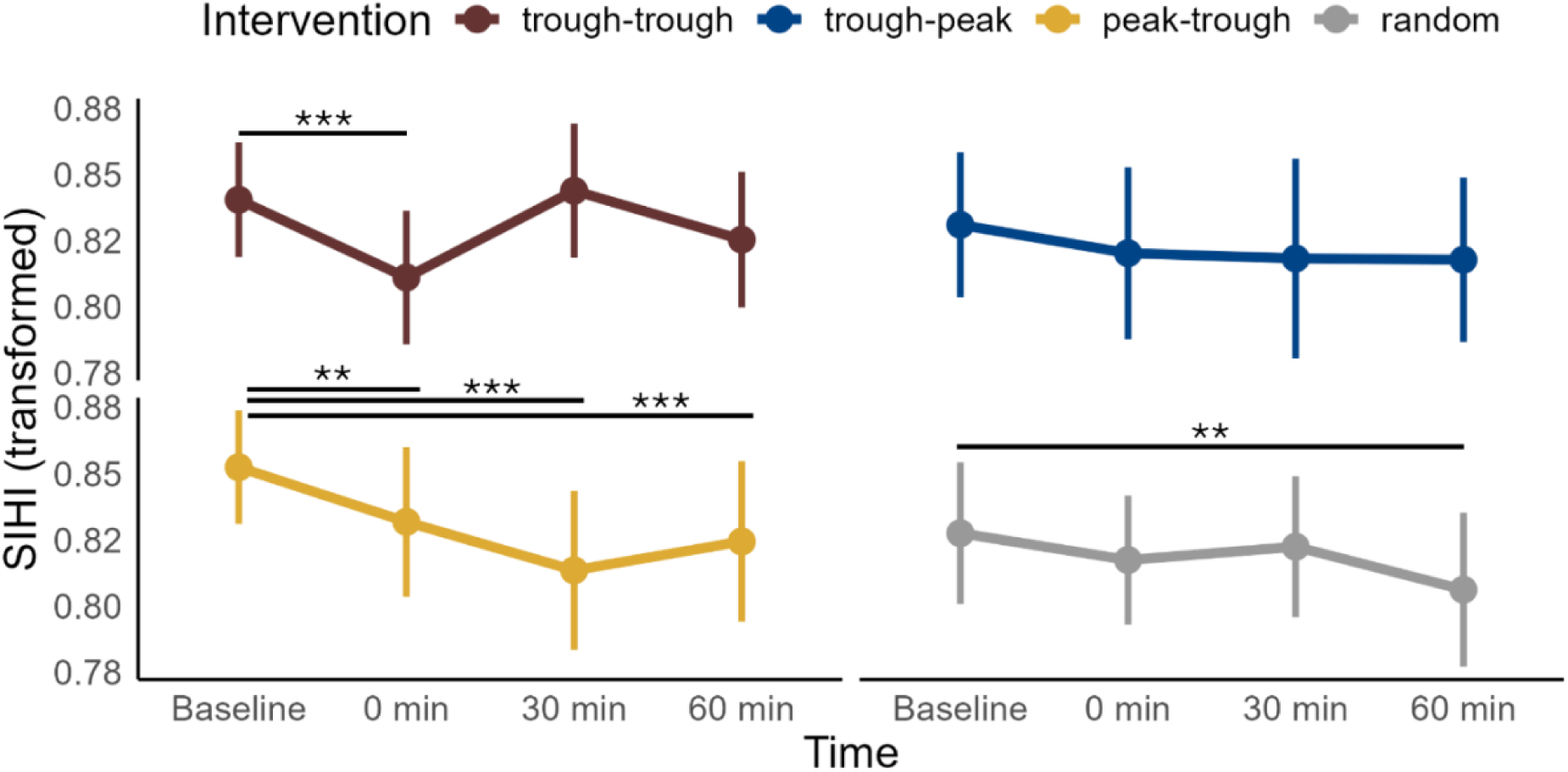
Effects of ccPAS intervention and time on SIHI. SIHI as a function of intervention type and recording time. Lower values indicate stronger inhibition. SIHI values were transformed using the fourth root. The plots display the means and standard errors of the mean (SEM) of all subjects (n = 16). SIHI values were pooled across all conditioning stimulus intensities, since **Intensity** had no significant triple interaction with **Intervention** and **Time**. Trough-positive peak intervention is labelled as “trough-peak”, positive peak-trough – as “peak-trough”, random phase – as “random”. Pairwise comparisons between post-intervention time points and the baseline were conducted for each intervention type using marginal means derived from the LME model, not the depicted data. Horizontal bars with asterisks indicate time points where the marginal mean differed significantly from the baseline. Significance codes: *** <0.001, ** <0.01, * <0.05.

Conditioned corticospinal excitability in the right test M1: For conditioned stimulation, a significant interaction was observed between **Intervention** and **Time** (χ² = 238, df = 9, p < 0.001, Fig. 3S), as well as between **Intervention** and **Intensity** (χ² = 10, df = 3, p < 0.05). Main effects were significant for **Intensity** (χ² = 4,935, df = 1, p < 0.001) and **Time** (χ² = 66, df = 3, p < 0.001). Across all intervention conditions, MEP amplitude decreased post-intervention, with the timing of these changes varying by condition (see Table 3S and Fig. 3S for pairwise comparisons). However, in the case of trough-trough ccPAS, MEP amplitude decreased at 0 min but then increased at 30 min compared to baseline. No significant differences were found between **Interventions** at any specific time point.

**Figure 3.**
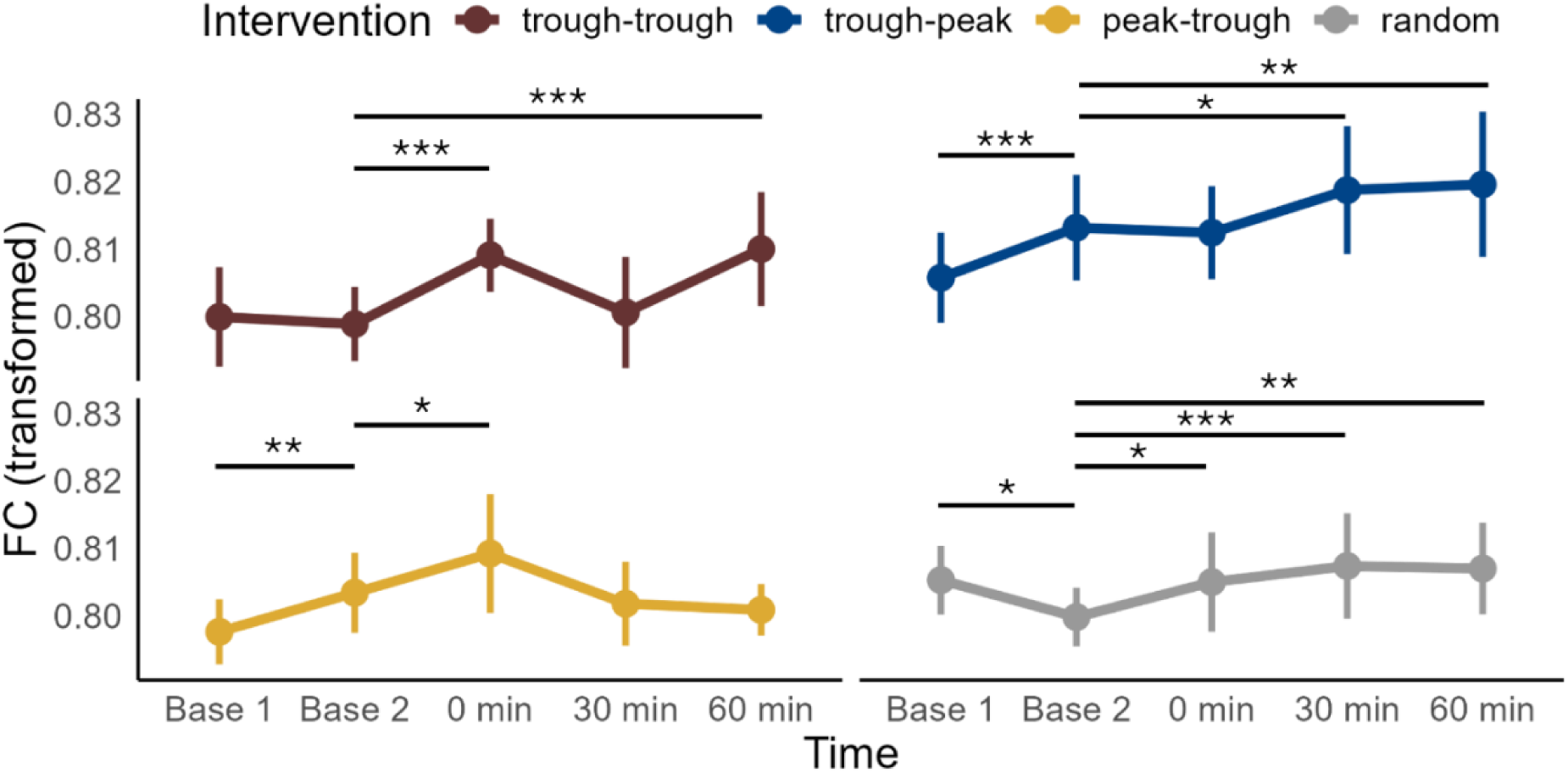
Effects of ccPAS intervention and time on wPLI. WPLI as a function of intervention type and recording time. Lower values indicate stronger inhibition. WPLI values were transformed using the fourth root. Higher values indicate stronger connectivity. The plots display the means and standard errors of the mean (SEM) of all subjects (n = 16). Trough-positive peak intervention is labelled as “trough-peak”, positive peak-trough – as “peak-trough”, random phase – as “random”. Pairwise comparisons between post-intervention time points and the baseline were conducted for each intervention type using marginal means derived from the LME model, not the depicted data. Horizontal bars with asterisks indicate time points where the marginal mean differed significantly from the baseline. Significance codes: *** <0.001, ** <0.01, * <0.05.

### Short-interval interhemispheric inhibition (SIHI)

A significant interaction was observed between **Intervention** and **Time** (χ² = 44, df = 9, p < 0.001, Fig. 2), as well as between **Intervention** and **Intensity** (χ² = 9, df = 3, p < 0.05, data not shown). Main effects were also significant for **Intensity** (χ² = 5,595, df = 1, p < 0.001) and **Time** (χ² = 44, df = 3, p < 0.001). Across all intervention conditions, except for the trough-positive peak condition, SIHI was strengthened post-intervention, with the timing of changes varying by condition (see Fig. 2 and Table 4S for pairwise comparisons). However, no significant differences were found between **Interventions** at any specific time point.

The changes in SIHI were not correlated with the average change (across trials, intervention conditions, and intensities) of MEP amplitude elicited by either unconditioned stimulation of the right test M1 or stimulation of the left conditioning M1 (Figs. 4SB,C). As expected, the changes in SIHI were correlated with the average change (across trials, intervention conditions, and intensities) of MEP amplitude elicited by conditioned stimulation of the right test M1 (Pearson’s r = 0.67, p < 0.001) (Fig. 4SD). This validated that SIHI measured what we assumed it to measure. We excluded the possibility that the observed changes in SIHI were driven by changes in unconditioned MEPs (i.e., corticospinal excitability) in the left conditioned M1.

### Functional connectivity (wPLI)

There was a significant interaction between **Intervention** and **Time** (χ² = 116, df = 12, p < 0.001, Fig. 3), as well as a significant main effect of **Time** (χ² = 97, df = 4, p < 0.001). Upon pairwise comparison, no intervention condition differed significantly from the random-phase condition at any time point. Meanwhile, functional connectivity increased post-intervention across all conditions, but the timing of this increase varied (see Fig. 3 and Table 5S for pairwise comparisons). Notably, all conditions except trough-trough showed significant differences between the two baseline measurements, indicating potential variability even in the absence of intervention. There was no correlation between the changes in SIHI and the average (across epochs and intervention conditions) functional connectivity indexed by wPLI (Fig. 4SA).

### Functional connectivity (phase lag shift)

We evaluated the effects of ccPAS intervention on functional M1-M1 connectivity expressed in shifting of the mean phase lag. Phase lag varied across **Subjects** (χ² = 40,129, df = 15, p < 0.001), yet was clustered around the population mean (Fig. 5S). No effect of **Intervention** was found on the shift in phase lag between pre- and post-intervention recordings. Notably, we did not explicitly test whether there was a non-zero shift in phase lag post-intervention but rather whether the (zero or non-zero) shift differed between **Intervention** types.

## Discussion

We showed here that M1-M1 ccPAS resulted in a significant strengthening of M1-M1 effective and functional connectivity, as measured by SIHI in dual-coil TMS (Fig. 2) and wPLI in resting-state EEG (Fig. 3), respectively. However, in contrast to our expectation, these changes were largely independent of the phase of the ongoing µ-rhythm in the conditioning and test M1, and were found also with random phase stimulation.

The strengthening in SIHI is also in contrast with the single other previous M1-M1 ccPAS study that tested SIHI before and after ccPAS and demonstrated a weakening in SIHI ^11^. The ccPAS interventions between the two studies are largely identical (i.e., left M1 served as conditioning M1, right M1 as test M1; 180 paired-pulses per intervention in the present study, 90 paired-pulses per intervention in the previous study; suprathreshold stimulus intensities for both pulses; low stimulation rate of ∼0.2 Hz in the present study and 0.05 Hz in the previous study; interstimulus interval included 10 ms in both studies; time points of post-ccPAS measurements of 0, 30 and 60 min in both studies). Significant differences were the use of smaller figure-of-eight TMS coils in the present study (outer diameter of each wing, 35 mm) compared to the previous study (70 mm) and a biphasic current waveform in the present study versus a monophasic waveform in the previous study. Significant differences in the induction of lasting changes in M1 excitability with biphasic versus monophasic repetitive TMS were observed in earlier studies ^27,28^. Therefore, we consider this the most likely reason for the strengthening vs. weakening of SIHI in the present vs. previous study^11^.

The ccPAS intervention led to an long-term potentiation-like increase in corticospinal excitability in the left conditioning M1 (Fig. S1), again in contrast to the previously observed effects, where no significant change in corticospinal excitability was found for the conditioning M1 ^11^. It is possible that the observed increased cortical excitability in the conditioning M1 has contributed to strengthening of SIHI in our study, as stronger conditioning pulses lead to stronger SIHI ^29,30^. This possibility is supported by previous studies that either applied low-frequency 1 Hz repetitive TMS to the left conditioning M1, resulting in a concomitant decrease of corticospinal excitability of the conditioning M1, and a weakening of SIHI ^31^, or quadripulse stimulation with an interpulse interval of 5 ms, resulting in a concomitant increase of corticospinal excitability of the conditioning M1, and a strengthening of SIHI ^32^.

The ccPAS intervention did not lead to a significant change (i.e., effect of Time) in corticospinal excitability of the right test M1, when measured with unconditioned test pulses (Fig. S2). This is where our results also diverge from the previous findings that showed a significant increase in corticospinal excitability of the test M1 ^11^. This increase in corticospinal excitability observed in the previous study may have contributed to the weakening of SIHI, as other previous studies demonstrated that SIHI weakened with increasing test MEP amplitudes ^4,33^. This was further supported by the observation that low-frequency 1 Hz repetitive TMS applied to the left conditioning M1 resulted in a concomitant increase in corticospinal excitability of the right test M1 and weakening of SIHI ^34^.

In order to assess the possible relevance of these contributions in altered corticospinal excitability of the conditioning M1 and the test M1 we performed correlations analyses with the ccPAS-induced change in SIHI. These did not show significant correlations of changes in SIHI with changes in MEP amplitude elicited from the left conditioning M1 or unconditioned right test M1 (Fig. 4SB,C), indicating that the changes in corticospinal excitability were of minor relevance for the observed ccPAS-induced strengthening of SIHI. A concomitant change in corticospinal excitability of the test M1 (increase) and SIHI (weakening) also occurred in the previous study ^11^, but the correlation of these changes was not examined. Pal and colleagues provided evidence for the independence of the changes in corticospinal excitability of the conditioning M1 and SIHI by correcting for the 1 Hz repetitive TMS-induced decrease in conditioning MEP amplitude ^31^. In accordance, the transcallosal projection neurons probably mediating SIHI are located in cortical layers II/III and are, therefore, a neuronal population distinct from the corticospinal neurons located in layer V, involved in MEP generation ^35^.

We have demonstrated previously that (1) corticospinal excitability is increased when TMS is delivered at the trough compared to the peak of the ongoing µ-rhythm ^15,36^ and (2) SIHI is stronger when both M1 are in synchrony with the phase of their ongoing µ-rhythms, compared to out of synchrony conditions ^17^. Based on these previous findings, we expected that brain state-dependent ccPAS will most effectively change SIHI in conditions of high M1-M1 effective connectivity (i.e., when the µ-rhythm of both M1 is in synchrony) and/or a high-excitability state of the conditioning M1 and/or the test M1 (i.e., when the µ-rhythm in the conditioning or test M1 is at trough). In accord with this expectation, a recent study demonstrated that bifocal in-phase transcranial alternating current stimulation (1.0 mA, 20 Hz, 20 min) over the bilateral sensorimotor cortices increased effective connectivity as measured with SIHI, but not functional connectivity indexed in the resting-state EEG by the imaginary part of coherency ^37^.

Why we did not find differential effects between the ccPAS conditions on the induced long-term increase in SIHI may have a number of reasons. SIHI as a marker of effective transcallosal connectivity is a net inhibition, consisting of a superimposition of strong interhemispheric inhibition and weaker interhemispheric facilitation ^38^. Both can change after intervention ^32^, and this renders unequivocal interpretation of changes in SIHI impossible. Furthermore, it is unclear if the different ccPAS interventions activated interhemispheric interactions in different ways. We have not directly tested SIHI during ccPAS. But our previous findings suggest that SIHI does not show a strong modulation with the phase (trough vs. positive peak) of the ongoing µ-rhythm ^17^, in contrast to a clear effect of µ-phase on corticospinal excitability ^15,17,36,39^. Finally, we targeted specifically the corticospinal neurons in layer V involved in the generation of MEP by employing a hot spot search (see Methods). However, the neurons mediating transcallosal interaction are distinct. They are located in layers II/III ^35^, and our recent advanced TMS mapping research has demonstrated that they are optimally activated some millimeters away from the hot spot but only suboptimally activated when targeting the hot spot with focal TMS (Vetter et al., in review). This suboptimal targeting also may have contributed to the failure in detecting significant differences in SIHI plasticity induction between the ccPAS interventions.

We observed a strengthening of functional connectivity across all ccPAS interventions, as indicated by the increase in wPLI in the resting-state EEG measurements (Fig. 3). Phase synchronization metrics, such as wPLI, identify statistical (i.e., undirected) associations characterized by stability of the phase lag between time series of electrophysiological recordings from spatially distinct brain areas. WPLI ^18^ minimizes the non-specific contribution of (almost-)zero-lag interactions, potentially caused by volume conduction and is, thus, expected to allow identifying true time-lagged functional couplings ^40^. Beyond the ubiquitous increase in phase lag stability, we observed no differential changes to the mean phase lag following the ccPAS interventions. While varying on population level, it was not susceptible to phase-dependent modulation. Nevertheless, the strengthening in functional connectivity did not correlate with the strengthening in SIHI (Fig. 4SA), suggesting independence of the underlying mechanisms.

## Conclusions

In summary, brain state-dependent ccPAS induces long-term changes in the transcallosal motor network. However, further research is needed to improve our understanding of the physiological underpinnings of this form of plasticity and optimize the efficacy of its induction by ccPAS or other interventions.

## Author contributions

M.E.: Conceptualization, Methodology, Investigation, Formal analysis, Writing— original draft preparation, Visualization. G.K.: Conceptualization, Methodology, Investigation, Software, Formal analysis, Writing—original draft preparation, Visualization. P.B.: Conceptualization, Formal analysis, Writing—review \& editing. U. Z.: Conceptualization, Supervision, Writing—review & editing, Funding acquisition.

## Data availability

Public sharing of raw data is not possible due to the data protection agreement with the participants. The raw data can be shared individually upon request to the corresponding author. The pre-processed data is available at: 10.5281/zenodo.15126575. The analysis code is available at: https://github.com/mariaermolova/M1M1PAS.

## Acknowledgements

This work was supported by the European Research Council (ERC Synergy) under the European Union’s Horizon 2020 research and innovation programme (ConnectToBrain; grant agreement No. 810377) (to U.Z.). G. Kozák declares salary support from sync2brain GmbH as a part-time employee, sync2brain GmbH is a University of Tübingen spin-off that commercializes a variant of the real-time EEG analysis device used in this research.

