## Supplementary Material for "Plasticity of interhemispheric motor cortex connectivity induced by brain state-dependent cortico-cortical paired-associative stimulation"

**Table 1S:** Pairwise comparisons of corticospinal excitability in the left conditioning M1 measured by MEP amplitude in the right FDI

Pairwise comparisons of marginal means from the LME model on the MEP amplitudes in the FDI of the right hand. Trough-positive peak intervention is labelled as “trough-peak”, positive peak-trough – as “peak-trough”, random phase – as “random”.

| Time | Intervention | contrast | estimate | SE | df | z.ratio | p.value |
|---|---|---|---|---|---|---|---|
| Pre | . | trough-trough - random | 0.079 | 0.215 | Inf | 0.366 | 0.779 |
| Pre | . | trough-peak - random | 0.336 | 0.215 | Inf | 1.563 | 0.236 |
| Pre | . | peak-trough - random | 0.277 | 0.215 | Inf | 1.287 | 0.318 |
| 0 | . | trough-trough - random | -0.119 | 0.215 | Inf | -0.552 | 0.729 |
| 0 | . | trough-peak - random | 0.249 | 0.215 | Inf | 1.157 | 0.371 |
| 0 | . | peak-trough - random | 0.325 | 0.215 | Inf | 1.509 | 0.242 |
| 30 | . | trough-trough - random | -0.166 | 0.215 | Inf | -0.773 | 0.620 |
| 30 | . | trough-peak - random | -0.103 | 0.215 | Inf | -0.478 | 0.729 |
| 30 | . | peak-trough - random | -0.055 | 0.215 | Inf | -0.254 | 0.834 |
| 60 | . | trough-trough - random | 0.038 | 0.215 | Inf | 0.176 | 0.860 |
| 60 | . | trough-peak - random | 0.277 | 0.215 | Inf | 1.286 | 0.318 |
| 60 | . | peak-trough - random | 0.101 | 0.215 | Inf | 0.470 | 0.729 |
| **.** | **random** | **0 - Pre** | **0.284** | **0.047** | **Inf** | **6.036** | **0.000** |
| **.** | **random** | **30 - Pre** | **0.463** | **0.047** | **Inf** | **9.837** | **0.000** |
| **.** | **random** | **60 - Pre** | **0.326** | **0.047** | **Inf** | **6.935** | **0.000** |
| . | trough-trough | 0 - Pre | 0.087 | 0.047 | Inf | 1.842 | 0.143 |
| **.** | **trough-trough** | **30 - Pre** | **0.218** | **0.047** | **Inf** | **4.631** | **0.000** |
| **.** | **trough-trough** | **60 - Pre** | **0.286** | **0.047** | **Inf** | **6.068** | **0.000** |
| **.** | **trough-peak** | **0 - Pre** | **0.197** | **0.047** | **Inf** | **4.178** | **0.000** |
| . | trough-peak | 30 - Pre | 0.024 | 0.047 | Inf | 0.512 | 0.729 |
| **.** | **trough-peak** | **60 - Pre** | **0.267** | **0.047** | **Inf** | **5.668** | **0.000** |
| **.** | **peak-trough** | **0 - Pre** | **0.332** | **0.047** | **Inf** | **7.048** | **0.000** |
| **.** | **peak-trough** | **30 - Pre** | **0.132** | **0.047** | **Inf** | **2.795** | **0.012** |
| **.** | **peak-trough** | **60 - Pre** | **0.151** | **0.047** | **Inf** | **3.202** | **0.004** |

**Figure 1S:** Effects of intervention on corticospinal excitability in the left conditioning M1 measured by MEP amplitude in the right FDI

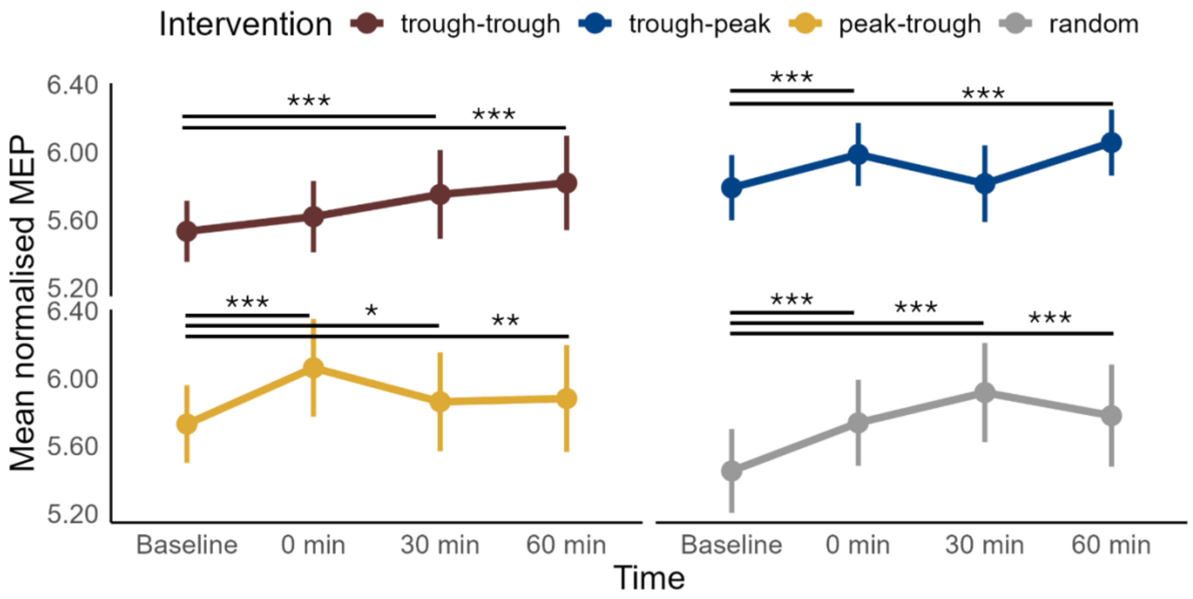


**Figure 1S.** Effects of ccPAS intervention and time on corticospinal excitability of the left conditioning M1 measured by MEP amplitude in the right FDI. MEP as a function of intervention type and recording time. Higher values indicate higher MEP amplitude. MEP values were transformed using the fourth root. The plots display means and standard errors of the mean (SEM) of all subjects (n = 16). MEP values were pooled across all conditioning stimulus intensities, since **Intensity** had no significant triple interaction with **Intervention** and **Time**. Trough-positive peak intervention is labelled as “trough-peak”, positive peak-trough – as “peak-trough”, random phase – as “random”. Pairwise comparisons between post-intervention time points and the baseline were conducted for each intervention type using marginal means derived from the LME model, not the depicted data. Horizontal bars with asterisks indicate time points where the marginal mean differed significantly from the baseline. Significance codes: *** <0.001, ** <0.01, * <0.05.

**Table 2S:** Pairwise comparisons of unconditioned corticospinal excitability in the right test M1 measured by MEP amplitude in the left FDI

Pairwise comparisons of marginal means from the LME model on the MEP amplitudes in the FDI of the left hand. Trough-positive peak intervention is labelled as “trough-peak”, positive peak-trough – as “peak-trough”, random phase – as “random”.

| Time | Intervention | contrast | estimate | SE | df | z.ratio | p.value |
|---|---|---|---|---|---|---|---|
| Pre | . | trough-trough - random | 0.188 | 0.191 | Inf | 0.981 | 0.582 |
| Pre | . | trough-peak - random | 0.070 | 0.191 | Inf | 0.366 | 0.807 |
| Pre | . | peak-trough - random | 0.304 | 0.191 | Inf | 1.589 | 0.336 |
| 0 | . | trough-trough - random | -0.237 | 0.191 | Inf | -1.237 | 0.573 |
| 0 | . | trough-peak - random | -0.158 | 0.191 | Inf | -0.824 | 0.614 |
| 0 | . | peak-trough - random | 0.183 | 0.191 | Inf | 0.955 | 0.582 |
| 30 | . | trough-trough - random | 0.330 | 0.191 | Inf | 1.727 | 0.289 |
| 30 | . | trough-peak - random | -0.048 | 0.191 | Inf | -0.251 | 0.837 |
| 30 | . | peak-trough - random | -0.007 | 0.191 | Inf | -0.038 | 0.970 |
| 60 | . | trough-trough - random | 0.144 | 0.191 | Inf | 0.756 | 0.635 |
| 60 | . | trough-peak - random | -0.225 | 0.191 | Inf | -1.178 | 0.573 |
| 60 | . | peak-trough - random | 0.067 | 0.191 | Inf | 0.353 | 0.807 |
| **.** | **random** | **0 - Pre** | **0.202** | **0.076** | **Inf** | **2.662** | **0.037** |
| . | random | 30 - Pre | 0.068 | 0.076 | Inf | 0.890 | 0.597 |
| . | random | 60 - Pre | 0.076 | 0.076 | Inf | 1.003 | 0.582 |
| **.** | **trough-trough** | **0 - Pre** | **-0.222** | **0.076** | **Inf** | **-2.917** | **0.032** |
| **.** | **trough-trough** | **30 - Pre** | **0.210** | **0.076** | **Inf** | **2.765** | **0.034** |
| . | trough-trough | 60 - Pre | 0.033 | 0.076 | Inf | 0.436 | 0.807 |
| . | trough-peak | 0 - Pre | -0.025 | 0.076 | Inf | -0.333 | 0.807 |
| . | trough-peak | 30 - Pre | -0.050 | 0.076 | Inf | -0.662 | 0.678 |
| **.** | **trough-peak** | **60 - Pre** | **-0.219** | **0.076** | **Inf** | **-2.882** | **0.032** |
| . | peak-trough | 0 - Pre | 0.081 | 0.076 | Inf | 1.070 | 0.582 |
| **.** | **peak-trough** | **30 - Pre** | **-0.243** | **0.076** | **Inf** | **-3.200** | **0.032** |
| . | peak-trough | 60 - Pre | -0.160 | 0.076 | Inf | -2.106 | 0.141 |

**Figure 2S:** Effects of intervention on unconditioned corticospinal excitability in the test right M1 measured by MEP amplitude in the left FDI

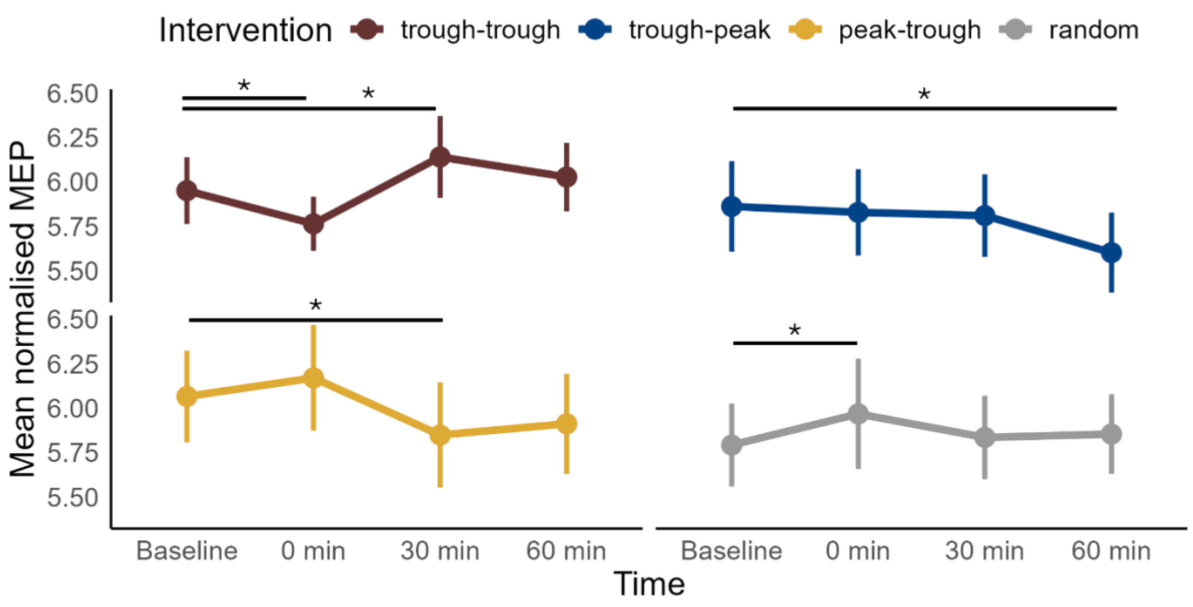


**Figure 2S.** Effects of ccPAS intervention and time on unconditioned corticospinal excitability in the right test M1 measured by MEP amplitude in the left FDI. MEP as a function of intervention type and recording time. Higher values indicate higher MEP amplitude. MEP values were transformed using the fourth root. The plots display the means and standard errors of the mean (SEM) of all subjects (n = 16). Trough-positive peak intervention is labelled as "trough-peak", positive peak-trough – as "peak-trough", random phase – as "random". Pairwise comparisons between post-intervention time points and the baseline were conducted for each intervention type using marginal means derived from the LME model, not the depicted data. Horizontal bars with asterisks indicate time points where the marginal mean differed significantly from the baseline. Significance codes: *** <0.001, ** <0.01, * <0.05.

**Table 3S:** Pairwise comparisons of conditioned corticospinal excitability in the right test M1 measured by MEP amplitude in the left FDI

Pairwise comparisons of marginal means from the LME model on the MEP amplitudes in the FDI of the left hand. Trough-positive peak intervention is labelled as “trough-peak”, positive peak-trough – as “peak-trough”, random phase – as “random”.

| Time | Intervention | contrast | estimate | SE | df | z.ratio | p.value |
|---|---|---|---|---|---|---|---|
| Pre | . | trough-trough - random | 0.140 | 0.208 | Inf | 0.670 | 0.635 |
| Pre | . | trough-peak - random | 0.019 | 0.208 | Inf | 0.090 | 0.928 |
| Pre | . | peak-trough - random | 0.303 | 0.208 | Inf | 1.452 | 0.293 |
| 0 | . | trough-trough - random | -0.257 | 0.208 | Inf | -1.234 | 0.372 |
| 0 | . | trough-peak - random | -0.133 | 0.208 | Inf | -0.638 | 0.635 |
| 0 | . | peak-trough - random | 0.164 | 0.208 | Inf | 0.789 | 0.608 |
| 30 | . | trough-trough - random | 0.341 | 0.208 | Inf | 1.637 | 0.244 |
| 30 | . | trough-peak - random | -0.080 | 0.208 | Inf | -0.385 | 0.764 |
| 30 | . | peak-trough - random | -0.061 | 0.208 | Inf | -0.294 | 0.802 |
| 60 | . | trough-trough - random | 0.209 | 0.208 | Inf | 1.004 | 0.473 |
| 60 | . | trough-peak - random | -0.080 | 0.208 | Inf | -0.385 | 0.764 |
| 60 | . | peak-trough - random | 0.131 | 0.208 | Inf | 0.630 | 0.635 |
| . | random | 0 - Pre | 0.066 | 0.043 | Inf | 1.552 | 0.263 |
| . | random | 30 - Pre | -0.054 | 0.043 | Inf | -1.278 | 0.372 |
| **.** | **random** | **60 - Pre** | **-0.118** | **0.043** | **Inf** | **-2.766** | **0.019** |
| **.** | **trough-trough** | **0 - Pre** | **-0.331** | **0.043** | **Inf** | **-7.762** | **0.000** |
| **.** | **trough-trough** | **30 - Pre** | **0.147** | **0.043** | **Inf** | **3.447** | **0.002** |
| . | trough-trough | 60 - Pre | -0.048 | 0.043 | Inf | -1.134 | 0.411 |
| . | trough-peak | 0 - Pre | -0.086 | 0.043 | Inf | -2.007 | 0.134 |
| **.** | **trough-peak** | **30 - Pre** | **-0.153** | **0.043** | **Inf** | **-3.600** | **0.002** |
| **.** | **trough-peak** | **60 - Pre** | **-0.217** | **0.043** | **Inf** | **-5.088** | **0.000** |
| . | peak-trough | 0 - Pre | -0.072 | 0.043 | Inf | -1.695 | 0.240 |
| **.** | **peak-trough** | **30 - Pre** | **-0.418** | **0.043** | **Inf** | **-9.817** | **0.000** |
| **.** | **peak-trough** | **60 - Pre** | **-0.289** | **0.043** | **Inf** | **-6.790** | **0.000** |

**Figure 3S:** Effects of intervention on conditioned corticospinal excitability in the test right M1 measured by MEP amplitude in the left FDI

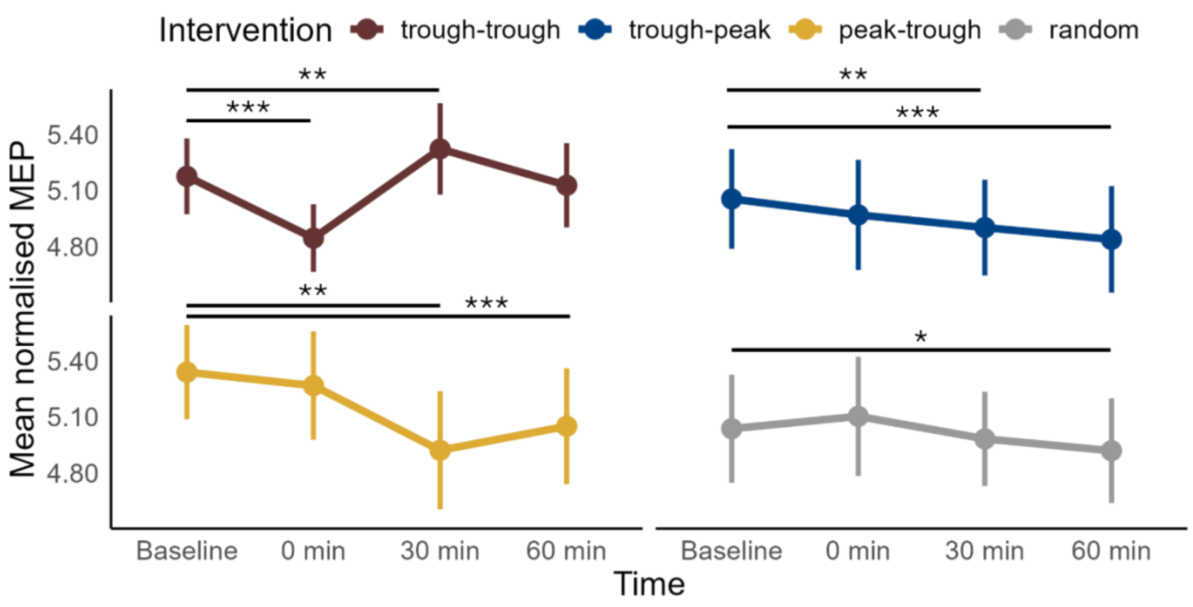


**Figure 3S.** Effects of ccPAS intervention and time on conditioned corticospinal excitability in the right test M1 measured by MEP amplitude in the left FDI. MEP as a function of intervention type and recording time. Higher values indicate higher MEP amplitude. MEP values were transformed using the fourth root. The plots display the means and standard errors of the mean (SEM) of all subjects (n = 16). MEP values were pooled across all conditioning stimulus intensities, since **Intensity** had no significant triple interaction with **Intervention** and **Time**. Trough-positive peak intervention is labelled as “trough-peak”, positive peak-trough – as “peak-trough”, random phase – as “random”. Pairwise comparisons between post-intervention time points and the baseline were conducted for each intervention type using marginal means derived from the LME model, not the depicted data. Horizontal bars with asterisks indicate time points where the marginal mean differed significantly from the baseline. Significance codes: *** <0.001, ** <0.01, * <0.05.

**Table 4S:** Pairwise comparisons of short-interval interhemispheric inhibition

Pairwise comparisons of marginal means from the LME model on SIHI. Trough-positive peak intervention is labelled as “trough-peak”, positive peak-trough – as “peak-trough”, random phase – as “random”.

| Time | Intervention | contrast | estimate | SE | df | z.ratio | p.value |
|---|---|---|---|---|---|---|---|
| Pre | . | trough-trough - random | 0.013 | 0.019 | Inf | 0.689 | 0.693 |
| Pre | . | trough-peak - random | 0.003 | 0.019 | Inf | 0.185 | 0.878 |
| Pre | . | peak-trough - random | 0.025 | 0.019 | Inf | 1.327 | 0.403 |
| 0 | . | trough-trough - random | -0.006 | 0.019 | Inf | -0.336 | 0.843 |
| 0 | . | trough-peak - random | 0.003 | 0.019 | Inf | 0.154 | 0.878 |
| 0 | . | peak-trough - random | 0.014 | 0.019 | Inf | 0.767 | 0.664 |
| 30 | . | trough-trough - random | 0.021 | 0.019 | Inf | 1.143 | 0.506 |
| 30 | . | trough-peak - random | -0.004 | 0.019 | Inf | -0.220 | 0.878 |
| 30 | . | peak-trough - random | -0.009 | 0.019 | Inf | -0.475 | 0.762 |
| 60 | . | trough-trough - random | 0.019 | 0.019 | Inf | 1.025 | 0.564 |
| 60 | . | trough-peak - random | 0.012 | 0.019 | Inf | 0.620 | 0.714 |
| 60 | . | peak-trough - random | 0.018 | 0.019 | Inf | 0.970 | 0.569 |
| . | random | 0 - Pre | -0.010 | 0.006 | Inf | -1.567 | 0.281 |
| . | random | 30 - Pre | -0.005 | 0.006 | Inf | -0.791 | 0.664 |
| **.** | **random** | **60 - Pre** | **-0.021** | **0.006** | **Inf** | **-3.303** | **0.006** |
| **.** | **trough-trough** | **0 - Pre** | **-0.029** | **0.006** | **Inf** | **-4.545** | **0.000** |
| . | trough-trough | 30 - Pre | 0.003 | 0.006 | Inf | 0.529 | 0.754 |
| . | trough-trough | 60 - Pre | -0.015 | 0.006 | Inf | -2.329 | 0.079 |
| . | trough-peak | 0 - Pre | -0.011 | 0.006 | Inf | -1.659 | 0.259 |
| . | trough-peak | 30 - Pre | -0.013 | 0.006 | Inf | -1.967 | 0.148 |
| . | trough-peak | 60 - Pre | -0.013 | 0.006 | Inf | -2.041 | 0.141 |
| **.** | **peak-trough** | **0 - Pre** | **-0.021** | **0.006** | **Inf** | **-3.193** | **0.007** |
| **.** | **peak-trough** | **30 - Pre** | **-0.039** | **0.006** | **Inf** | **-6.025** | **0.000** |
| **.** | **peak-trough** | **60 - Pre** | **-0.028** | **0.006** | **Inf** | **-4.340** | **0.000** |

**Table 5S:** Pairwise comparisons of weighted phase lag index

Pairwise comparisons of marginal means from the LME model on wPLI. Trough-positive peak intervention is labelled as “trough-peak”, positive peak-trough – as “peak-trough”, random phase – as “random”.

| Time | Intervention | contrast | estimate | SE | df | z.ratio | p.value |
|---|---|---|---|---|---|---|---|
| Pre 1 | . | trough-trough - random | -0.005 | 0.006 | Inf | -0.901 | 0.518 |
| Pre 1 | . | trough-peak - random | 0.000 | 0.006 | Inf | 0.080 | 0.942 |
| Pre 1 | . | peak-trough - random | -0.008 | 0.006 | Inf | -1.291 | 0.359 |
| Pre 2 | . | trough-trough - random | 0.000 | 0.006 | Inf | -0.073 | 0.942 |
| Pre 2 | . | trough-peak - random | 0.013 | 0.006 | Inf | 2.216 | 0.064 |
| Pre 2 | . | peak-trough - random | 0.004 | 0.006 | Inf | 0.659 | 0.632 |
| 0 | . | trough-trough - random | 0.005 | 0.006 | Inf | 0.837 | 0.543 |
| 0 | . | trough-peak - random | 0.008 | 0.006 | Inf | 1.299 | 0.359 |
| 0 | . | peak-trough - random | 0.005 | 0.006 | Inf | 0.769 | 0.571 |
| 30 | . | trough-trough - random | -0.007 | 0.006 | Inf | -1.258 | 0.359 |
| 30 | . | trough-peak - random | 0.011 | 0.006 | Inf | 1.792 | 0.162 |
| 30 | . | peak-trough - random | -0.006 | 0.006 | Inf | -0.952 | 0.503 |
| 60 | . | trough-trough - random | 0.003 | 0.006 | Inf | 0.534 | 0.707 |
| 60 | . | trough-peak - random | 0.013 | 0.006 | Inf | 2.267 | 0.060 |
| 60 | . | peak-trough - random | -0.006 | 0.006 | Inf | -0.989 | 0.500 |
| **.** | **random** | **Pre 1 - Pre 2** | **0.005** | **0.002** | **Inf** | **2.830** | **0.016** |
| **.** | **random** | **0 - Pre 2** | **0.005** | **0.002** | **Inf** | **2.378** | **0.049** |
| **.** | **random** | **30 - Pre 2** | **0.008** | **0.002** | **Inf** | **4.045** | **0.000** |
| **.** | **random** | **60 - Pre 2** | **0.007** | **0.002** | **Inf** | **3.614** | **0.002** |
| . | trough-trough | Pre 1 - Pre 2 | 0.000 | 0.002 | Inf | 0.185 | 0.912 |
| **.** | **trough-trough** | **0 - Pre 2** | **0.010** | **0.002** | **Inf** | **5.377** | **0.000** |
| . | trough-trough | 30 - Pre 2 | 0.001 | 0.002 | Inf | 0.320 | 0.830 |
| **.** | **trough-trough** | **60 - Pre 2** | **0.010** | **0.002** | **Inf** | **5.678** | **0.000** |
| **.** | **trough-peak** | **Pre 1 - Pre 2** | **-0.007** | **0.002** | **Inf** | **-4.087** | **0.000** |
| . | trough-peak | 0 - Pre 2 | -0.001 | 0.002 | Inf | -0.456 | 0.744 |
| **.** | **trough-peak** | **30 - Pre 2** | **0.005** | **0.002** | **Inf** | **2.722** | **0.020** |
| **.** | **trough-peak** | **60 - Pre 2** | **0.007** | **0.002** | **Inf** | **3.846** | **0.001** |
| **.** | **peak-trough** | **Pre 1 - Pre 2** | **-0.006** | **0.002** | **Inf** | **-3.488** | **0.002** |
| **.** | **peak-trough** | **0 - Pre 2** | **0.005** | **0.002** | **Inf** | **2.856** | **0.016** |
| . | peak-trough | 30 - Pre 2 | -0.002 | 0.002 | Inf | -1.051 | 0.478 |
| . | peak-trough | 60 - Pre 2 | -0.003 | 0.002 | Inf | -1.608 | 0.223 |

**Figure 4S:** Correlation analyses of ccPAS-induced changes in SIHI vs. functional connectivity and MEP amplitudes

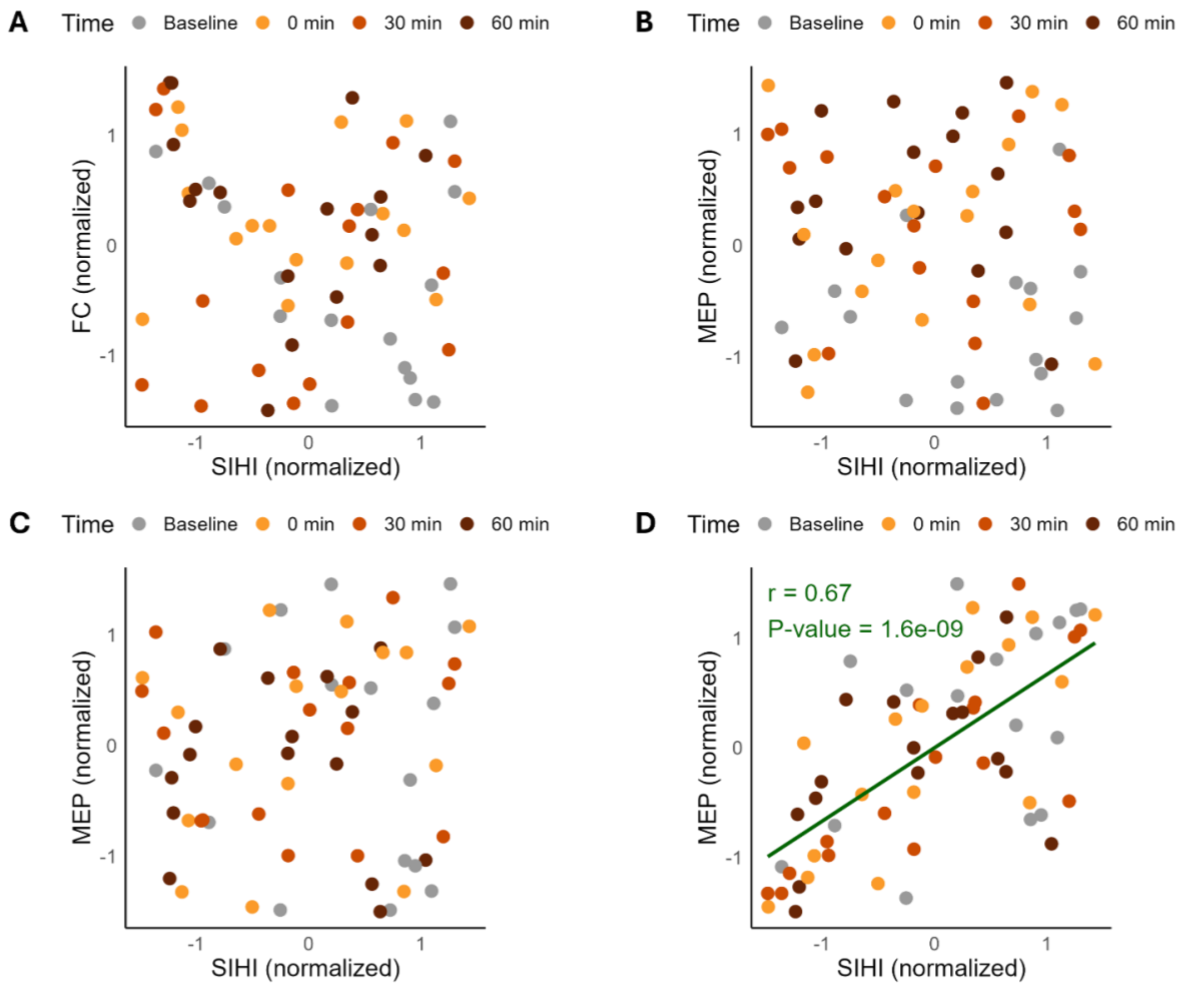


**Figure 4S.** Correlation analyses of ccPAS-induced changes in SICI vs. functional connectivity and MEP amplitudes. Data were averaged across individual trials or epochs, intervention conditions, and, where applicable, stimulation intensities. Each point represents one time point (relative to intervention) from one subject. Points are color-coded by time of measurement. Data were transformed as for the LME models and subsequently normalized via z-score across time points within each subject. Pearson's correlation analysis (two-sided) was used to assess correlation. Green line indicates the linear fit wherever correlation was significant. **A**: Correlation between SIHI and functional connectivity (FC), measured by wPLI. The first baseline was removed from FC data to match SIHI. **B**: Correlation between SIHI and corticospinal excitability of the left conditioning M1, measured by MEP amplitude from the right FDI. **C**: Correlation between SIHI and corticospinal excitability of the right test M1, measured by MEP amplitude from the left FDI during unconditioned (single-pulse) stimulation. **D**: Correlation between SIHI and inhibition of the right test M1, measured by MEP amplitude from the left FDI during conditioned (paired-pulse) stimulation.

**Figure 5S:** Distribution of M1-M1 phase lags at the population level

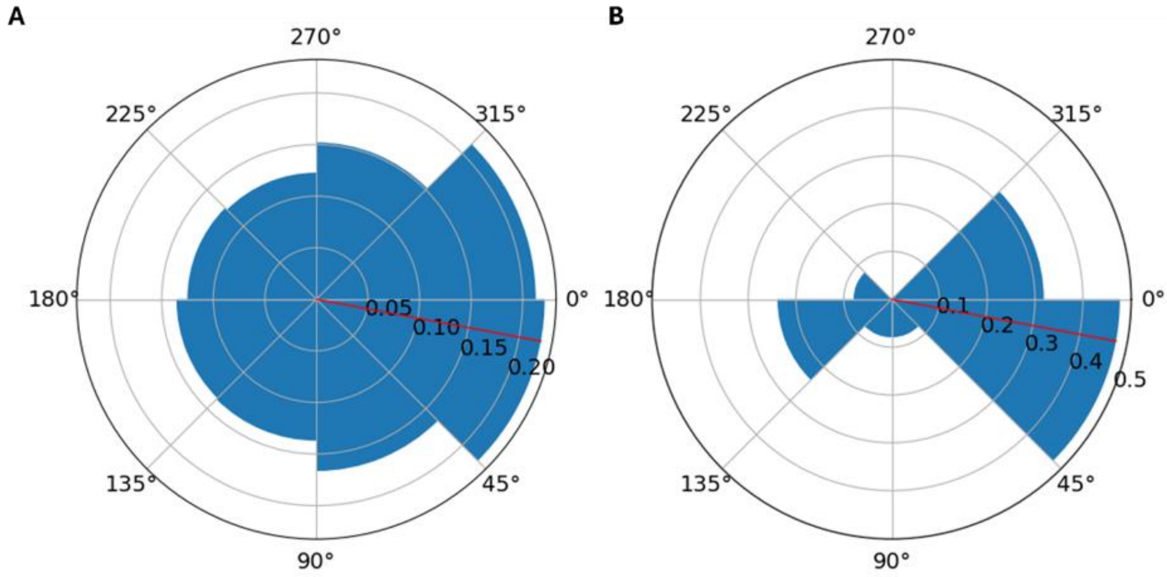


**Figure 5S.** Distribution of M1-M1 phase lags at the population level. **(A)** Distribution of individual phase lags (n = 1,117,000) between the µ-rhythms of the right and left M1 in the resting state, pooled from all subjects. The red bar represents mean of the distribution (11° ± 1.95° SD). The phase lags were calculated as $\varphi_{right} - \varphi_{left}$, i.e., the oscillation in the right M1 assumed to be the preceding one. **(B)** Distribution of subject-means of phase lags between the right and left M1 (n = 16). The red bar represents the mean of the population distribution of individual phase lags (same as in **(A),** for reference). Phase lag varied significantly across participants but was clustered around the population mean: 56% of participants featured a mean phase lag within a ±22.5° range around the population mean.
